# A growth-inhibitory dUTPase from *Mosigvirus* AV110 disrupts cell division and host physiology in *Escherichia coli*

**DOI:** 10.64898/2026.10.06.757002

**Authors:** Saar S.F. Van Overfelt, Viktor Hundtofte Mebus, Jonathan Brewer, Rob Lavigne, Anders Nørgaard Sørensen, Lone Brøndsted

**Author notes:** For correspondence: Lone Brøndsted, Department of Veterinary and Animal Sciences, University of Copenhagen, Stigbøjlen 4, 1870 Frederiksberg C, Denmark.

## Abstract

dUTPases are essential enzymes for nucleotide metabolism as they catalyze the hydrolysis of dUTP to dUMP, thus preventing the toxic incorporation of uracil into DNA. dUTPases are found in all domains of life, including bacteriophages, where they serve multiple functions to support phage propagation. Phage-encoded dUTPases reduce cellular dUTP levels and provide precursors for *de novo* dTTP synthesis and the synthesis of modified nucleotides, which are important to avoid DNA-based phage defense systems. In this study, we focused on the dimeric dUTPase Gp135, encoded by the *Mosigvirus* phage AV110, and examined its effect on growth, cell morphology, DNA replication and cell division of AV110’s host, extended-spectrum β-lactamase-producing *E. coli*. We observed that Gp135 significantly inhibited growth by disrupting DNA replication and cell division. This resulted in cytoplasmic voids, cell elongation, and ultimately cell death. We further demonstrated the diversity of phage-encoded dUTP/dCTPases through phylogenetic analysis, identifying four distinct clusters: one comprising the most common dUTPases, which form trimers; two more divergent clusters comprising dimeric dUTP/dCTPases; and one new cluster of more diverse, uncharacterized dUTPases and dNTPases. Overall, our results highlight the evolutionary and functional diversity of phage-encoded dUTP/dCTPases, with the *Mosigvirus* AV110 dUTP/dCTPase illustrating their role in supporting phage propagation.

## 1. Introduction

Enzymes like dUTPases play an essential role in nucleotide metabolism by catalyzing the hydrolysis of dUTP to dUMP [1]. By hydrolyzing dUTP to dUMP, dUTPases reduce intracellular dUTP levels and promote the incorporation of thymine, rather than the structurally similar uracil, into DNA. This is important to prevent mutations and maintain DNA integrity [1]. They are present across all kingdoms of life, including mammals [2], fungi [3], bacteria [4], viruses [5], and also viruses that infect bacteria, known as bacteriophages (phages) [6,7], although not every organism encodes a dUTPase [4]. dUTPases are highly diverse in sequence [8] and are classified into two evolutionarily distinct groups, with one forming a trimer or monomer [1], and the other forming a dimer or tetramer [9]. These two enzyme classes share no sequence similarity, and the sequence motifs critical for their activity vary [8].

During phage infection, dUTPases play a crucial role in regulating the nucleotide pools within the host cell [1]. This regulation is essential to provide sufficient nucleotides, especially dTTP, to support phage genome replication. To ensure adequate dTTP levels, it must be synthesized *de novo*, using dUMP as a precursor. During the infection process, dUMP is produced by the deamination of dCTP to dUTP, which is subsequently hydrolyzed by a dUTPase to dUMP [1]. Alternatively, some dimeric phage-encoded dUTPases, such as those encoded by the *Staphylococcus* phages 55 and phiNM1 and the *Escherichia* phages T2 and T4, can hydrolyze dCTP to dCMP[6,10–12]. This is then followed by the deamination of dCMP to dUMP. This dUMP is subsequently converted to dTMP by a thymidylate synthase and phosphorylated to dTTP [1]. The necessary enzymes for synthesizing dTTP from dCTP (dUTPase, dCMP deaminase, thymidylate synthase, and deoxynucleotide monophosphate kinase) are typically encoded by the phage. Bacteria also exploit the need of phages to have sufficient nucleotides available to defend themselves against phage infection, for example, by reducing dCTP or dGTP levels through a dCTP deaminase or dGTPase, respectively [13]. The presence of phage T7 mutants that can overcome the dCTP deaminase defense [13] and a counter-defense system encoded by phage T7 that inhibits the host dGTPase [14] highlights the importance of maintaining the nucleotide pools within the cell for phage infection.

Another function of dUTPases during phage infection is to counteract host defense mechanisms. Bacteria encode many defense systems, including the aforementioned dCTP deaminase and dGTPase, as well as restriction-modification and CRISPR-Cas systems, to protect against phage infection [15]. Conversely, phages have evolved to overcome those defense systems, for example, by incorporating (hyper)modified nucleotides into their genome to evade certain restriction-modification and CRIPSR-Cas systems [16,17]. Examples of such modifications include 5-glucosyl-hydroxymethylcytosine in *Tequatrovirus* [18] and 5-arabinosyl-arabinosyl-hydroxycytosine in the related *Winklervirus* and *Mosigvirus* [17]. It was shown that *Mosigvirus* phages, which encode a dimeric dUTPase that also uses dCTP as a substrate, catalyze the first step in the formation of those modified cytosines: the hydrolysis of dCTP to dCMP. This is followed by the addition of a hydroxyl group by dCMP hydroxylase, yielding 5-hydroxy-dCMP. This modified nucleotide is subsequently phosphorylated by a phage- and a host-encoded kinase to form 5-hydroxy-dCTP, which is incorporated into the DNA by the DNA polymerase. After being incorporated into the DNA, 5-hydroxycytosine is hypermodified through the addition of two arabinose molecules by a glycosyl transferase and an arabinose-5-ara-hydroxycytosine transferase [17,19].

The protein Gp135, encoded by the *Mosigvirus* phage AV110, was previously identified as a homolog of the T4 dUTP/dCTPase and was shown to inhibit the growth of the ESBL *E. coli* strain ESBL098 [20]. Our primary objective was to gain a deeper understanding of the impact of Gp135 on the cell morphology, DNA replication and cell division of ESBL098. Next, we aimed to address the unknown prevalence and diversity of dimeric and trimeric dUTPases encoded by phages by determining the phylogenetic distribution of these enzymes in relation to their taxonomic classification. Our findings highlight the evolutionary and functional diversity of phage-encoded dUTP/dCTPases using *Mosigvirus* AV110 Gp135 to demonstrate the impact on host physiology when redirecting host resources toward phage propagation.

## 2. Materials and Methods

### Bacterial strains and plasmids

The bacterial strains and plasmids used in this study are shown in Table S1. Bacteria were grown in Lysogeny Broth (LB) (BD 240110) and LB agar (BD 240230), when necessary supplemented with 50 µg/ml kanamycin, and grown overnight at 37°C with shaking at 180 rpm, unless otherwise stated.

### Growth inhibition assay

A diverse set of electrocompetent ESBL/AmpC *E. coli* strains (Table S1) was transformed with pPS26::*gp135* by electroporation. A growth inhibition assay was performed in biological triplicates by preparing a tenfold dilution series of an overnight culture of each strain in LB with 50 µg/ml kanamycin. Next, 10 µl of each dilution was spotted onto LA plates containing kanamycin, with and without 5 mM rhamnose. The plates were incubated overnight at 37°C, after which the growth-inhibitory effect was assessed by determining the log reduction.

### Growth curves

Growth curves were obtained in biological triplicates by growing ESBL098 pPS26::*gp135* to an OD_600_ of 0.2 in LB with kanamycin. Next, 100 µl of the culture was added to 100 µl of LB with kanamycin, with or without rhamnose, at a final concentration of 5 mM in a 96-well plate. The growth curve was obtained by measuring OD_600_ every 15 min for 24 h using the PowerWave XS microplate reader (BioTek) and Gen5 software.

### Time-lapse microscopy with and without live-dead staining

Time-lapse microscopy with and without live-dead staining was performed using ESBL098 transformed with pPS26::*gp135* or the pPS26 empty vector. An overnight culture of each strain was diluted 1/100 in fresh LB supplemented with kanamycin and grown for 1 h at 37°C with shaking at 120 rpm. Then, the culture was diluted to an OD_600_ of 0.01, and 1 µl of the culture was spotted on a 1.2% agarose pad in LB with 50 µg/ml kanamycin, 5 mM rhamnose, and, when necessary, 1 µM propidium iodide and sealed with a coverslip. Time-lapse microscopy was performed on three colonies of each strain, using an inverted Olympus IX83 microscope and a CellVivo incubator module set at 37°C. Phase-contrast images were acquired every 5 min for 25 h using a 1.4 N. A. 100x oil-immersion objective. When propidium iodide was added, fluorescence images were additionally acquired every 10 minutes. Image analysis was performed in Fiji [21].

### Fluorescence microscopy

An overnight culture of ESBL098 transformed with pPS26::*gp135* or the empty plasmid was diluted to an OD_600_ of 0.05 in fresh LB with kanamycin and incubated at 37°C with shaking at 120 rpm. After 30 min, 5 mM rhamnose and 10 µg/ml Nile Red were added to the culture, and the culture was incubated again at 37°C with shaking at 120 rpm. After 1 h, 2 h, and 4 h incubation in the presence of rhamnose, 100 µM OGDA, and 5 µg/ml DAPI were added, and the culture was briefly incubated. Next, cells were pelleted, and this pellet was resuspended in 1 ml 4.6% formaldehyde for overnight incubation at 4°C. The next day, the fixed cells were washed and resuspended in PBS. A 1.5 µl aliquot of the fixed cells was spotted on a 1.2% agarose pad in PBS, and a coverslip was applied. Images were acquired with an inverted Olympus IX83 microscope using a 1.4 N. A. 100x oil-immersion objective. Phase-contrast and fluorescence images were acquired using the U-FMCHE (Nile Red), U-FBNA (OGDA), and U-FUNA (DAPI) filter cubes. Fifteen images were acquired for each sample and time point. The images were processed and analyzed in Fiji [21].

### Phylogenetic analysis of phage-encoded dUTPases

The amino acid sequences of dUTPases, dCTPases, dUTP pyrophosphatases, and dCTP pyrophosphatases encoded by *Caudoviricetes* were downloaded from NCBI using the RefSeq database (retrieved on 4/7/2025), and one sequence for each genus was included. Sequences from phages with no assigned taxonomy were removed. The remaining 168 sequences, together with the sequence of four known dimeric dUTPases (NP_049646, AAK66980, YP_873980, and YP_240523), two known trimeric dUTPases (YP_006959 and ABF71603), and Gp135 from phage AV110 (WPK33974), were aligned using CLC Main Workbench 25.0 with the *E. coli* K12 thymidylate synthase (NP_417304) as an outgroup. A phylogenetic tree was constructed from the alignment using the Neighbor Joining algorithm, the Jukes-Cantor protein distance measurements, and 100 replicates for bootstrapping. The resulting phylogenetic tree was visualized in iTOL [22].

## 3. Results

### The AV110 dUTPase affects the growth of ESBL *E. coli*

In a screen of early-expressed proteins of *Mosigvirus* phage AV110, we identified Gp135, which encodes a dUTPase, as a protein with a growth-inhibitory effect on ESBL098 [20]. The AV110 dUTPase caused at least a four-log reduction in the growth of ESBL098 (Fig.1). To investigate its growth-inhibitory impact in a broader range of strains, the protein was expressed in a diverse panel of ESBL/AmpC *E. coli* strains [23] and *E. coli* Stellar™. This demonstrated that the growth-inhibitory effect was not limited to ESBL098 but was observed in all tested strains (Fig.1). The growth-inhibitory effect on plates ranged from almost complete inhibition in ESBL049 to an approximate three-log reduction in ESBL054. In all tested strains, smaller colonies were visible, likely due to resistance to dUTPase, potentially conferred by mutations in the expression vector. Insertions in the expression vector causing resistance against the phage protein have been observed previously in a similar assay [20]. The broad-spectrum effect of expressing the AV110 dUTPase within (ESBL/AmpC) *E. coli* suggests interference with a conserved process essential for growth.

**Fig. 1:**
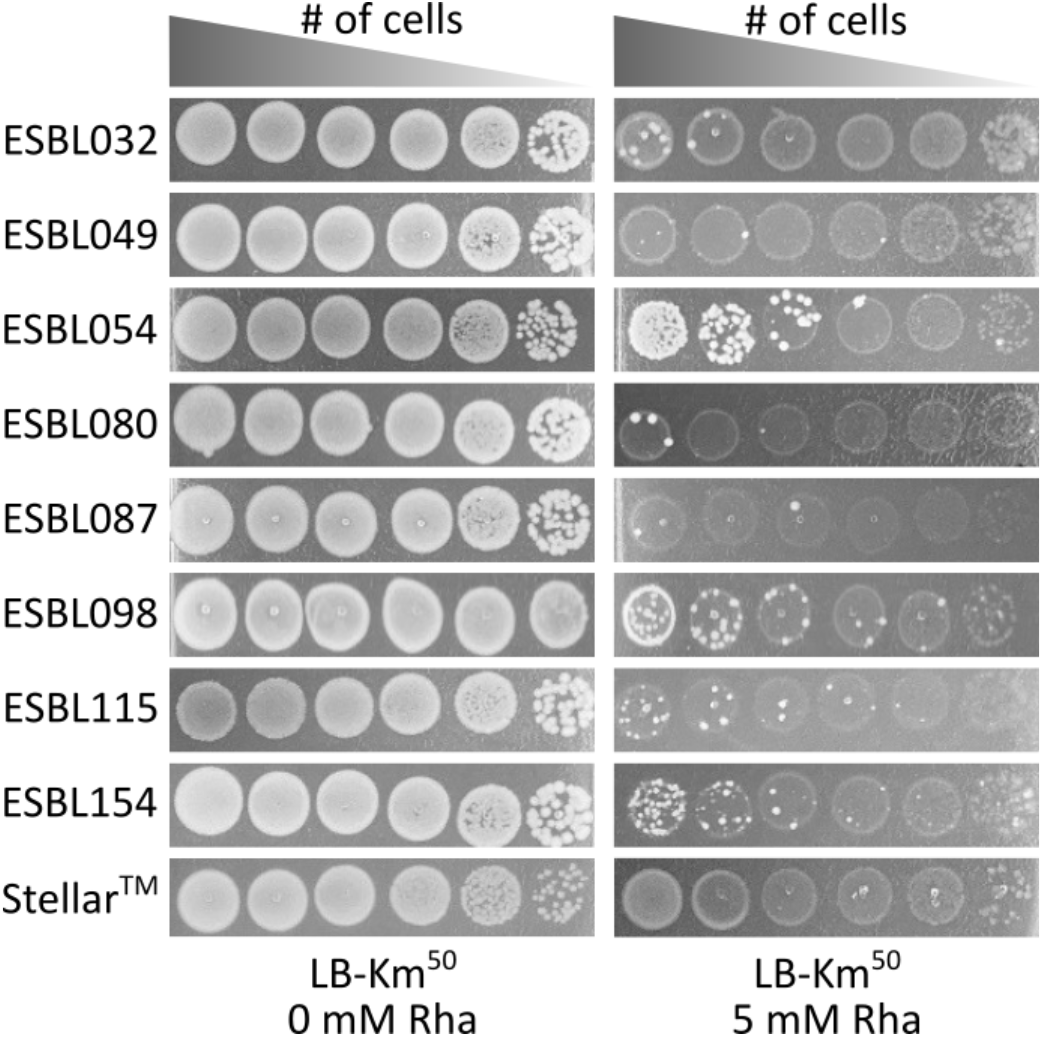
The AV110 dUTPase inhibits the growth of diverse *E. coli* strains. Tenfold dilutions of diverse ESBL *E. coli* strains and Stellar™ transformed with pPS26::*gp135* were spotted on LB agar plates with (right) and without (left) rhamnose induction.

### AV110 dUTPase influences cell morphology and cellular processes

To investigate the effect of the dUTPase on the growth of ESBL/AmpC *E. coli* over time, the growth of ESBL098 with recombinant expression of the dUTPase was followed in liquid medium through OD_600_ measurements. Expression of the dUTPase did not impact growth during the first two hours, likely due to the time needed to produce sufficient protein to affect the cells. However, between two and nine hours post-induction, a decrease in OD_600_ was observed, followed again by an increase (Fig. 2A). Time-lapse microscopy suggested that this OD_600_ increase may be attributed to the formation of elongated cells (Fig. 2C and Movie S1). A live-dead stain indicated that the dUTPase expression eventually led to cell death and significantly more cell death was observed compared to the empty vector control (Fig. 2B and Movie S2).

**Fig. 2:**
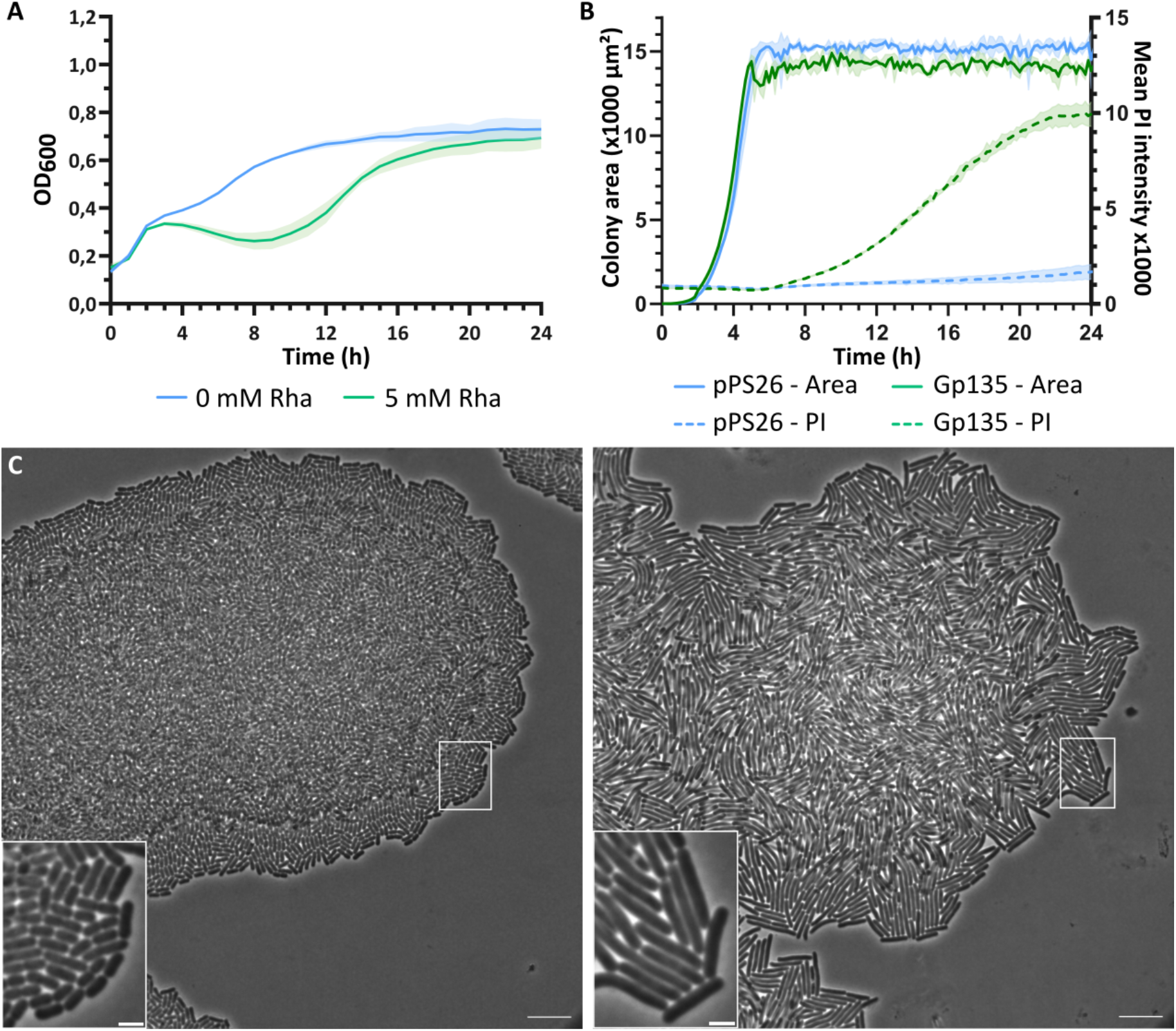
Expression of the dUTPase changes ESBL098 cell morphology. (A) The growth of ESBL098 pPS26::*gp135* was followed in liquid LB in the presence (green) and absence (blue) of 5 mM rhamnose. The mean of the measured OD_600_ and the standard deviation of three biological replicates are shown. (B) The detected colony size and the average propidium iodide (PI) intensity over time during time-lapse microscopy with live-dead staining of ESBL098 transformed with the pPS26 empty vector and pPS26::*gp135* are shown. The average and standard deviation of three biological replicates are shown. (C) Phase contrast image of ESBL098 transformed with the pPS26 empty vector (left) and pPS26::*gp135* (right) after growth for four hours in the presence of 5 mM rhamnose. Scale bar = 10 µm, scale bar insert = 2 µm.

To better understand how the dUTPase impacts cellular processes, fluorescence microscopy was performed one, two, and four hours after induction, staining the cell membrane, sites of active peptidoglycan synthesis, and DNA. After one hour of dUTPase expression, no differences were observed in any of the three visualized structures compared to the control (Fig. S1). However, after two hours, no septum formation was observed, suggesting that cells were no longer dividing. Additionally, the DNA signal intensity varied between cells, and the DNA did not appear to be replicated, as only one DNA signal was visible in each cell, whereas multiple signals would be expected if DNA had been replicated and segregated (Fig. 3A). These observations became more pronounced after four hours of induction (Fig. 3B). Peptidoglycan was still actively being synthesized in the cell wall, but without the initiation of septum formation, and DNA intensity varied between cells with no visible signs of replication. Unlike the images after one and two hours, differences were observed in the phase-contrast image and the cell membrane. In the phase contrast image, bright bands were visible in a number of cells along the cell and at the poles, while the cell membrane, DNA, and peptidoglycan synthesis remained intact at these locations (Fig. 3B). The bands spanned the width of the cell and suggest a region of lower optical density, possibly due to a cytoplasmic void. Foci were also visible in the cell membrane, indicating localized accumulations or disruptions of the cell membrane, although this is likely associated with cell death rather than a direct effect of the dUTPase (Fig. 3B). These results demonstrate that the AV110 dUTPase expression hinders DNA replication, leading to the inhibition of cell division, cell elongation, and cell death.

**Fig. 3:**
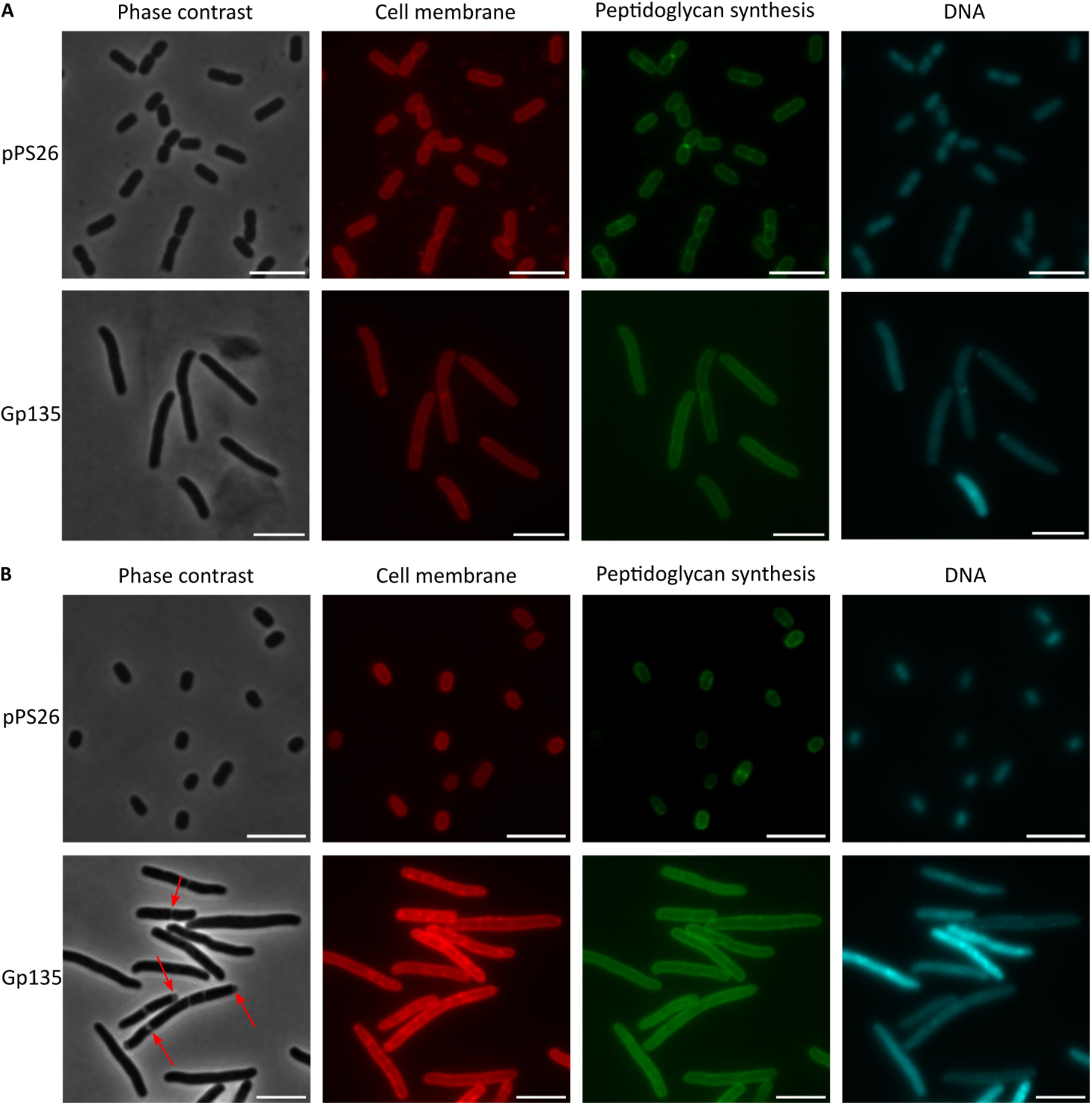
Expression of AV110 dUTPase changes cellular processes of ESBL098. Phase contrast and fluorescence microscopy images with staining of the cell membrane, sites with active peptidoglycan synthesis, and DNA, respectively, of ESBL098 transformed with the empty vector pPS26 or pPS26::*gp135* after (A) two hours and (B) four hours of induction. The red arrows indicate cytoplasmic voids along the longitudinal axis of the cell and at the cell poles. Representative images from fifteen images per condition per time point are shown. Scale bar = 5 µm.

### AV110 dUTPase is a homolog of the T4 dUTP/dCTPase

Gp135 was previously annotated as a nucleoside triphosphate pyrophosphohydrolase and was identified as an early-expressed protein based on its homology to the T4 dUTP/dCTPase [20]. Using an InterPro [24] and NCBI conserved domain search [25,26], a conserved dUTPase domain (EC 3.6.1.23) was identified, which belongs to the all-α NTP pyrophosphatase superfamily (EC 3.6.1.8). dUTPases within this family typically form a homodimer [27], but two homodimers can further interact to form a tetramer [10]. Structural predictions with AlphaFold 3 [28] showed that the AV110 dUTPase is composed exclusively of alpha helices (Fig. 4A) and likely interacts to form a dimer or tetramer (ipTM score = 0.88 and 0.90, respectively). The predicted tetrameric structure of the AV110 dUTPase is highly similar to the predicted tetrameric structures of the T2 and T4 dUTP/dCTPase (Fig. 4A; T2 RMSD = 0.738, T4 RMSD = 3.168). Also at the sequence level, the AV110 dUTPase has significant similarity with the T2 and T4 dUTP/dCTPase, and the critical residues in the different sequence motifs that are important for forming the ligand binding site and binding with Mg2+, as determined by Moroz and colleagues [27], are conserved between the three proteins (Fig. 4B). These results indicate that the AV110 dUTPase likely functions as a tetrameric dUTP/dCTPase that upon overexpression inhibits DNA replication in *E. coli* by depleting dUTP and dCTP.

**Fig. 4:**
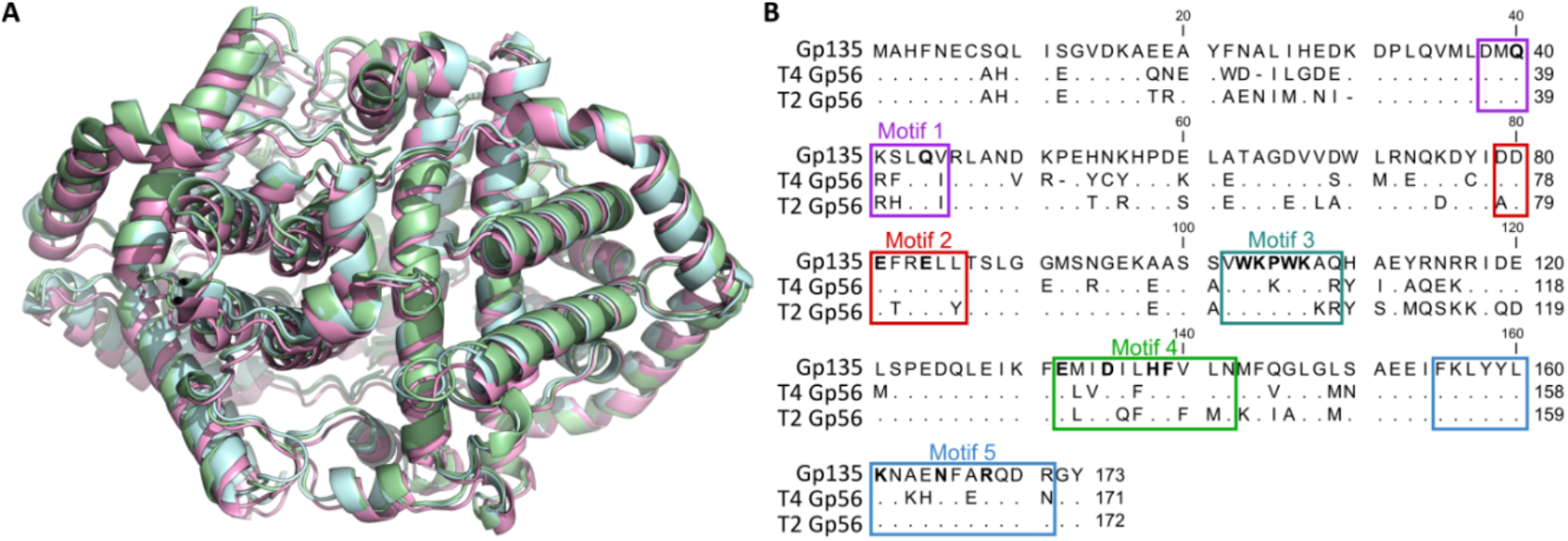
The AV110 dUTPase shows structural and sequence similarity to the dUTP/dCTPases from phages T2 and T4. (A) Overlay of the tetrameric protein structure of the AV110 dUTPase (light cyan), the T4 dUTPase (pink; RMSD = 3.168), and the T2 dUTPase (green; RMSD = 0.738). All protein structures were predicted as tetramers with AlphaFold 3. (B) Alignment of the amino acid sequence of the AV110 dUTPase (Gp135) and the dUTP/dCTPase from phages T4 and T2, encoded by *gp56*. Identical residues are indicated with dots. The conserved sequence motifs are indicated, and the critical residues in each motif according to Moroz and colleagues [27] are shown in bold.

### Phage-encoded dUTP/dCTPases form four distinct clusters

We showed that the AV110 dUTPase has a broad effect on the host, suggesting that it plays a central role in phage-host interactions and phage DNA replication. To determine whether these functions are conserved across phages, a phylogenetic tree of phage-encoded dUTP/dCTPases was constructed. The sequence of one dUTP/dCTPase per phage genus was selected from the NCBI RefSeq database, resulting in 168 protein sequences from 17 phage families and 34 subfamilies. The majority of these proteins are annotated as dUTPase, with a small number of proteins annotated as dCTPase or nucleoside triphosphate pyrophosphohydrolase (dNTPase). As references, the sequences of the AV110 dUTPase, four known dimeric dUTPases from *Staphylococcus* phages 55 and phiNM1, and *Escherichia* phages T2 and T4 [6,9,10,12,29], along with two trimeric dUTPases from *Staphylococcus* phage 80α and *Escherichia* phage T5 [30,31], were included. The resulting phylogenetic tree revealed distinct clades, indicating different evolutionary origins of dimeric and trimeric dUTPases (Fig. 5). Trimeric dUTPases formed a single cluster of closely related proteins, all annotated as dUTPase, while dimeric dUTPases grouped into two separate clusters also containing proteins annotated as dCTPase and dNTPase. Sequence alignment of the dimeric dUTPase clusters showed little to no similarity between the two clusters. The dimeric dUTPases from *Staphylococcus* phages 55 and φNM1 clustered together, whereas those from *Escherichia* phages T2, T4, and AV110 formed the second group. The first cluster also included dNTPases from *Bacillus* phages as well as dUTPases encoded by *Staphylococcus* phages. The second cluster primarily consisted of dUTPases encoded by phages from the *Straboviridae* family, which infect *E. coli, Acinetobacter, Pseudomonas*, and *Aeromonas*, among other organisms. The close evolutionary relatedness between the dUTP/dCTPases in this cluster with the AV110, T2 and T4 dUTPase indicates that the enzymes in this cluster likely all function as dUTPase and dCTPase, with similar effects on their host. Notably, some *Straboviridae*-encoded dUTPases were also found within the trimeric dUTPase cluster.

**Fig. 5:**
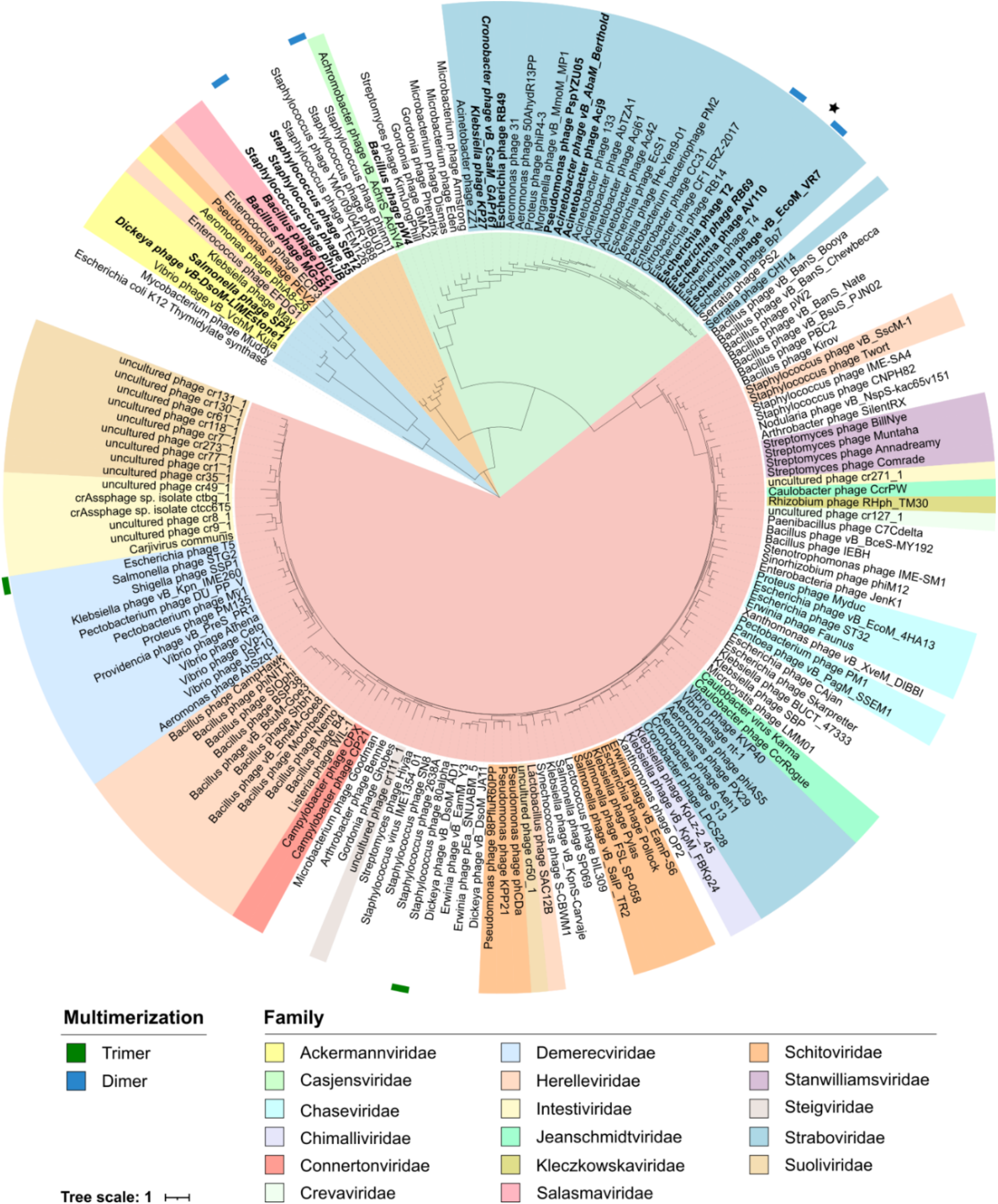
Phage-encoded dUTP/dCTPases form four distinct phylogenetic clusters. The sequences of dUTP/dCTPases encoded by *Caudoviricetes* were extracted from NCBI using the RefSeq database and one sequence of each genus was included. Thymidylate synthase encoded by *E. coli* K12 was added as an outgroup and used to root the tree. The sequence of known trimeric and dimeric dUTPases was added and those are indicated in the outer ring in green and blue, respectively. The dUTPase of AV110 is indicated with a star. The background colors of the labels represent the taxonomic classification at the family level. The branches of the tree are colored according to the different clusters of dUTP/dCTPases: one cluster of trimeric dUTPases (red), two clusters of dimeric dUTPases (orange and green), and the cluster with dUTPases that have not been studied (blue). Genes annotated as dCTPase are indicated in bold, while genes annotated as nucleoside triphosphate pyrophosphohydrolase are indicated in bold-italic.

A fourth cluster was formed by dUTPases and dNTPases encoded by *Ackermannviridae, Herelleviridae*, and *Schitoviridae* phages, although none of those proteins have been studied. Additionally, there was great diversity within this cluster, represented by low bootstrap support (20%), which suggests that this cluster could be further divided into two subclusters with more robust bootstrap support (71% and 66%) and the dUTPase from phage Muddy as a distinct lineage. Structural predictions of some of those sequences further showed the diversity within this cluster (Fig. S2). Some dUTPases had an all-alpha structure, partially resembling that of dimeric dUTPases (Fig. S2 A-B), while others had some structural similarity to trimeric dUTPases (Fig. S2 C-D). Together, these results highlight the great diversity within phage-encoded dUTP/dCTPases, with a fourth cluster of yet unknown enzymes.

## 4. Discussion

dUTPases are common in all types of living organisms, including phages [1]. In *Mosigvirus* AV110, this is encoded by Gp135, despite being annotated as a dNTPase. The effect of the dUTPase on AV110’s host, ESBL *E. coli*, was further investigated through its impact on multiple cellular processes upon overexpression. Overexpression of AV110 dUTPase inhibited growth across a diverse range of (ESBL/AmpC) *E. coli* strains. We hypothesize that this is due to the depletion of dCTP from the nucleotide pools, which is essential for host DNA replication. This halted DNA replication, leading to disruption of cell division, cell elongation, and, ultimately, cell death. After four hours of dUTPase expression, cytoplasmic voids were observed along the cell and at the poles, resembling those observed after three hours of thymine starvation in *E. coli*, which causes thymine-less death [32]. Similarly to AV110 dUTPase expression, thymine starvation additionally caused cell elongation and widening but resulted in a more condensed nucleoid in the middle of the cell than what we observed. The difference in nucleoid condensation could be due to the complete absence of dTTP upon thymine starvation, while AV110 dUTPase expression likely only partially depletes the nucleotide pool. Upon depletion of dGTP or dCTP from the nucleotide pools, similar observations have been made, including inhibition of cell growth and filament formation, and impaired DNA replication [33,34]. This suggests that the phenotype caused by AV110 dUTPase expression could be rescued by adding the various nucleotides. An analysis of the nucleotide pools in the cell upon AV110 dUTPase expression could further confirm its activity as a dUTP/dCTPase. However, these findings illustrate how phage-encoded dUTP/dCTPases, such as the AV110 dUTPase, can disrupt host cellular processes.

It is expected that the AV110 dUTPase contributes to various functions carried out by dUTP/dCTPases during phage infection. The essentiality of T4 dUTPase underscores the importance of these distinct functions [35]. First, because of the dUTPase domain and its similarity to the T2 and T4 dUTPases, the AV110 dUTPase likely hydrolyzes dUTP, which is crucial for preventing the incorporation of uracil into DNA and maintaining DNA integrity. Secondly, because T2 and T4 dUTPases can also hydrolyze dCTP, we hypothesize that the AV110 dUTPase also hydrolyzes dCTP. This activity is essential for the formation of 5-arabinosyl-arabinosyl-hydroxycytosine (5ara-ara-hC) by catalyzing the first step in the formation of this modified cytosine. Recent findings demonstrate that phage AV110 also encodes the other genes necessary for converting dCMP to 5ara-ara-hC, which helps the phage to evade DNA-based host defense systems [17]. Thirdly, the potential dUTP/dCTPase activity of the AV110 dUTPase is essential for regulating the nucleotide pools in the cells, as it provides dUMP and dCMP, both precursors for *de novo* synthesis of dTTP. Thereby, the AV110 dUTPase activity contributes to the successful replication of the phage genome.

The close phylogenetic relationship between the AV110 dUTPase and dUTP/dCTPases primarily encoded by *Straboviridae* phages suggests that the dimeric structure and function of the AV110 dUTPase is conserved in this cluster. It is therefore likely that the enzymes in this cluster share the ability to interfere with the host nucleotide pools, resulting in inhibited DNA replication and cell division and causing cell death. This may represent a widespread strategy of phages to modulate its host and promote phage replication. As previously observed, a second, distinct cluster of dimeric dUTPases encoded by phages infecting *Staphylococcus* and *Bacillus* was identified [27]. Although these clusters are evolutionarily distant, dUTPases in both groups exhibit dCTPase activity in addition to dUTPase activity [10–12], raising the possibility of either a shared ancestral origin with subsequent divergence or convergent evolution leading to the independent emergence of dCTPase activity. In both scenarios, the repeated emergence of dCTPase activity suggests an important selective advantage of extended hydrolyzing activity during infection. It additionally suggests that phage-encoded dUTP/dCTPases from both clusters cause similar effects on their host.

In contrast, trimeric dUTPases form a single, relatively conserved cluster and appear to be most prevalent in phages, consistent with observations in other organisms [27]. However, no trimeric phage-encoded dUTPases have yet been reported with extended hydrolytic activity. Given that most bacterial hosts already encode their own dUTPase, it could be expected that these enzymes have a more limited effect on their host. The identification of a fourth, more diverse cluster of putative dUTP/dNTPases, primarily encoded by *Ackermannviridae* phages indicate a gap in our current understanding of phage-encoded nucleotide metabolism. The inclusion of additional sequences, along with functional characterization of these enzymes would be beneficial to offer more conclusive insights on their evolutionary relatedness and biological role. Looking back at the entire phylogenetic tree, it was observed that several phage families encode both trimeric and dimeric dUTPases. This may reflect horizontal gene transfer from the host genome or from integrated prophages, as shown for eukaryotic viruses and their hosts [36] and may suggest a more flexible acquisition of nucleotide metabolism genes in response to selective pressures.

In conclusion, we showed that overexpression of the *Mosigvirus* AV110 dUTP/dCTPase in *E. coli* resulted in cell death by inhibiting DNA replication. It is hypothesized that this hydrolyzing activity contributes to successful phage infection by maintaining DNA integrity, providing defense against restriction-modification systems, and regulating the cellular nucleotide pools. Lastly, there is greater diversity of phage-encoded dUTP/dCTPases, with the presence of a fourth cluster of yet-unknown enzymes.

## General

During the preparation of this work, the authors used Copilot to refine the writing. After using this tool, the authors reviewed and edited the content as needed and take full responsibility for the publication’s content.

## Author contributions

S.S.F. Van Overfelt: Conceptualization, Formal analysis, Investigation, Methodology, Visualization, Writing – original draft, Writing – review & editing

V. H. Mebus: Investigation, Methodology, Writing – review & editing

J. Brewer: Conceptualization, Writing – review & editing

R. Lavigne: Conceptualization, Funding acquisition, Writing – review & editing

A. N. Sørensen: Conceptualization, Supervision, Writing – review & editing

L. Brøndsted: Conceptualization, Funding acquisition, Project administration, Supervision, Writing – review & editing

## Data Availability

Not applicable

