## Supplementary data for "A growth-inhibitory dUTPase from *Mosigvirus* AV110 disrupts cell division and host physiology in *Escherichia coli*"

### Supplementary Materials

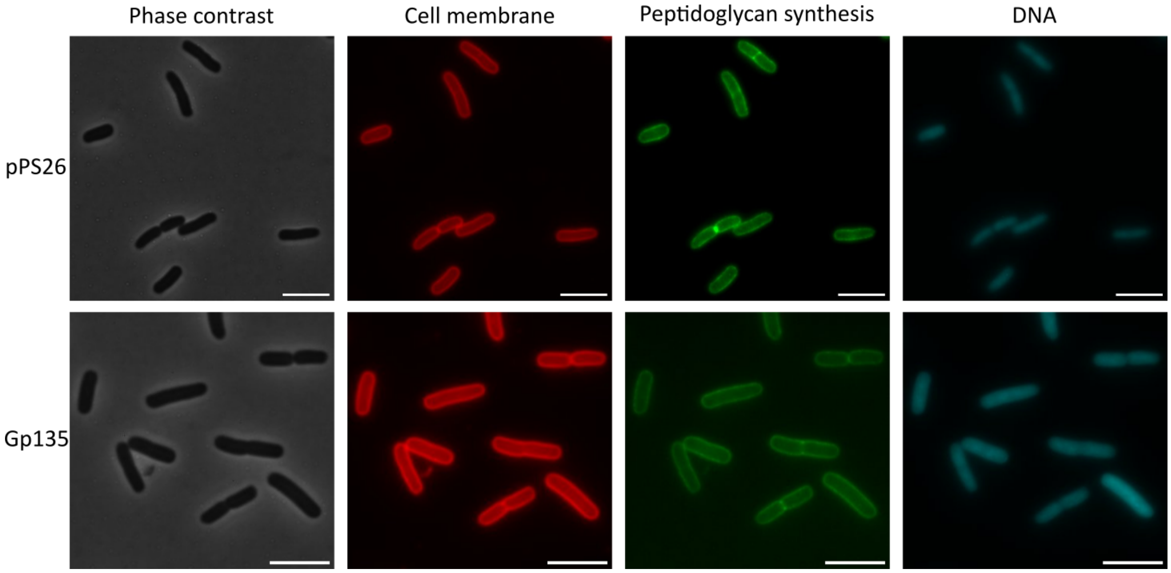


Fig. S1: Phase contrast and fluorescent microscopy with staining of the cell membrane, peptidoglycan synthesis, and DNA of ESBL098 transformed with the empty construct or pPS26::*gp135* and grown for one hour in the presence of 5 mM rhamnose. Representative images of 15 images per condition are shown. Scale bar = 5 µm.

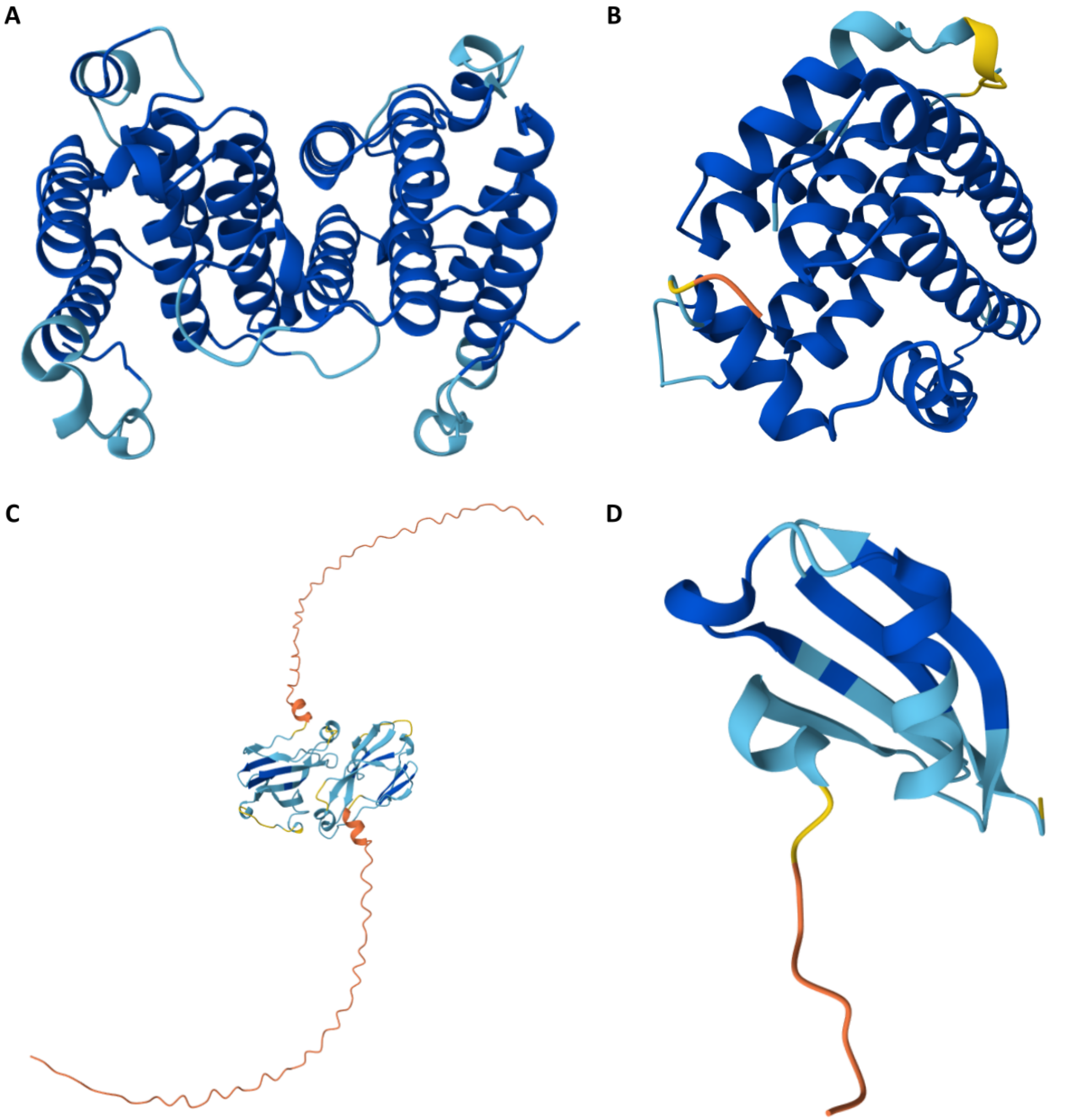


Fig. S2: Protein structure predicted with AlphaFold 3 of the dUTPase/dNTPase encoded by (A) *Salmonella* phage SP1, (B) *Aeromonas* phage phiA8-29, (C) *Pseudomonas* phage PEV2, and (D) *Enterococcus* phage ECP3.

Table S1: List of the bacterial strains and plasmids used in this study.

| **Bacteria** | **Source/Reference** |
|---|---|
| ESBL032 | [23] |
| ESBL049 | [23] |
| ESBL054 | [23] |
| ESBL080 | [23] |
| ESBL087 | [23] |
| ESBL115 | [23] |
| ESBL154 | [23] |
| ESBL098 pPS26 | [20] |
| ESBL098 pPS26::*gp135* | [20] |
| *E. coli* Stellar$^{TM}$ pPS26::*gp135* | [20] |
| **Plasmids** | |
| pPS26::*gp135* | [20] |

Movie S1: Time-lapse microscopy of ESBL098 pPS26::*gp135* grown for 24 h in the presence of 5 mM rhamnose.

Movie S2: Time-lapse microscopy of ESBL098 pPS26::*gp135* grown for 27 h in the presence of 5 mM rhamnose and 1 µM propidium iodide.
